# The spatial correspondence and genetic influence of inter-hemispheric connectivity with white matter microstructure

**DOI:** 10.1101/325787

**Authors:** Jeroen Mollink, Stephen M. Smith, Lloyd T. Elliott, Michiel Kleinnijenhuis, Marlies Hiemstra, Fidel Alfaro-Almagro, Jonathan Marchini, Anne-Marie van Cappellen van Walsum, Saad Jbabdi, Karla L. Miller

**Author notes:** Equal contribution (Saad Jbabdi, Karla L. Miller).

## Abstract

Microscopic features (i.e., microstructure) of axons affect neural circuit activity through characteristics such as conduction speed. Deeper understanding of structure-function relationships and translating this into human neuroscience has been limited by the paucity of studies relating axonal microstructure in white matter pathways to functional connectivity (synchrony) between macroscopic brain regions. Using magnetic resonance imaging data in 11354 subjects, we constructed multi-variate models that predict the functional connectivity of pairs of brain regions from the microstructural signature of white matter pathways that connect them. Microstructure-derived models provide predictions of functional connectivity that were significant in up to 86% of the brain region pairs considered. These relationships are specific to the relevant white matter pathway and have high reproducibility. The microstructure-function relationships are associated to genetic variants (single-nucleotide polymorphisms), co-located with genes *DAAM1* and *LPAR1*, that have previously been reported to play a role in neural development. Our results demonstrate that variation in white matter microstructure across individuals consistently and specifically predicts functional connectivity, and that this relationship is underpinned by genetic variability.

## Introduction

Communication between brain regions is achieved by axons grouped in white matter pathways, and properties of these axonal (structural) connections are highly relevant to brain function (e.g., functional connectivity). However, it is not simply the presence or absence of a connection, but also the tissue architecture (i.e., microstructure) of white matter that influences brain function. For example, axonal diameter, myelination and length all affect the precise timing of neural signals, which is crucial to synchronizing network dynamics^1^.

Much of our knowledge about structural connectivity in the brain comes from animals^2^, human lesions^3^, and post-mortem human dissections^4^. These approaches have relatively large biological specificity and interpretability but are limited in their ability to characterize inter-individual differences. More recently, diffusion MRI (dMRI) has emerged as a powerful *in vivo* tool for studying the brain’s structural connections^5^. Although limited in spatial resolution^6^, dMRI has the unique ability to estimate the trajectories of white matter bundles (i.e., tractography) as well as some microstructural properties of these bundles, through models linking the within-voxel dMRI signal to tissue architecture. An important benefit of dMRI is that it enables us to characterize inter-individual differences, even in large cohorts (e.g., UK Biobank^7^). dMRI thus has the potential to relate individual variations in white matter microstructure to differences in brain function, which can also be characterized with MRI.

Diffusion MRI has already been used to investigate structure – function relationships, mostly relating the presence and connectivity of a white matter tract to the functional coupling between the regions it is connecting^8–12^. Importantly, these studies relate the macroscopic organization of the network to brain function but did not aim to establish whether the microstructure of a white matter tract relates to the functional communication it establishes between brain areas.

A few studies have demonstrated the potential for dMRI to establish relationships between microstructure and function. For instance, the commonly-used metric fractional anisotropy (FA) is a measure of diffusion directionality that is biologically non-specific, being sensitive to many properties including axon density, size and myelination^13^. Mean FA in the cingulum bundle has been shown to correlate with functional connectivity between the medial frontal cortex and the posterior cingulate cortex, but not with functional connectivity derived from elsewhere in the brain^14^. Similar associations were found in callosal motor fibres connecting the hand areas in both hemispheres^15^. However, these studies focus on the tract connecting a single pair of regions and summarise a tract’s microstructure with a single quantity (e.g. FA averaged over the entire tract).

In this work, we address whether functional connectivity between brain regions is mediated by microstructure of white matter pathways that connect them. Unlike previous literature, we generate models that capture rich spatial representation of a tract’s microstructural profile (i.e., the microstructural signature). We show that these models can predict functional connectivity and demonstrate that the microstructure-function link is a general and reproducible principle in the human brain. In addition to diffusion tensor-based metrics, several microstructural measures were quantified using Neurite Orientation Dispersion and Density Imaging (NODDI^16^), a more sophisticated dMRI biophysical model. Measures derived from the NODDI model aim to provide greater biological specificity than diffusion tensor-model parameters such as FA. We consider interhemispheric connectivity between pairs of homotopic regions (i.e. the homologous region in the two cerebral hemispheres) that are connected by commissural white matter axons that run through the corpus callosum. We build a set of regression models to relate the tract’s microstructural profile to functional connectivity for a large number of paired homotopic regions. To test specificity of these models, we additionally built control models linking functional connectivity between a homotopic pair to tracts that do not connect them.

The models linking white matter microstructural signature to functional connectivity were trained on data from a large cohort of subjects (n=7481 subjects) and then were applied to an out-of-sample validation cohort (n=3873 subjects) in the UK Biobank project^7^. This dataset has the further benefit of enabling us to investigate what genetic variants underpin the relationship between microstructure and function of the human brain using a genome-wide association study (GWAS)^17^. Using the regression models described above, we identified a set of single-nucleotide polymorphisms (SNPs) that are significantly associated with functionally relevant microstructure in the brain^18^. The identified SNPs are co-located with genes that have been reported to play an important role in axonal guidance and cortical development, suggesting that microstructure-function relationships may be shaped in early development.

## Results

In our primary analysis, we tested for microstructure-function relationships between homotopic brain regions and the callosal pathways connecting them using dMRI and resting-state functional MRI (fMRI) data from subjects in the UK Biobank project^7^. All subjects were selected based upon usable resting-state fMRI and dMRI data, in addition to genetic inclusion criteria (see Methods section). The activity of homotopic region pairs is often synchronized, with high functional connectivity^19,20^. We focus on the corpus callosum in this study, because it is well defined and relatively immune to partial volume effects compared to other pathways.

### Functional connectivity

We previously conducted a group-average decomposition of resting-state fMRI data using Independent Components Analysis (ICA), which yielded 55 components corresponding to resting-state networks^7^. For the work here, more finely-grained functional “nodes” were then generated from these components by first splitting each component into its constituent parts for right and left hemispheres, and further splitting if a component still contained non-contiguous brain areas. Homologous regions for the two hemispheres were then identified as nodes with high symmetric similarity, producing 81 homotopic pairs (see Fig. 1.A). Functional connectivity was estimated at the single-subject level by partial correlation of the average BOLD signal time-series (equivalent to regressing out the time-series from all other regions prior to calculating pairwise correlations). The resulting connectivity matrix is given in Fig. 1.B as the mean partial correlation across all subjects. Entries in this matrix are ordered first by hemisphere and then by region number, such that inter-hemispheric connections are given in the upper-right and lower-left quadrants. Inter-hemispheric homotopic connections, shown on the diagonals of these quadrants, express on average the strongest connections in the brain, larger than intra-hemispheric or heterotopic inter-hemispheric connections (see Fig 1.C), in agreement with previous studies^19,20^.

**Figure 1.**
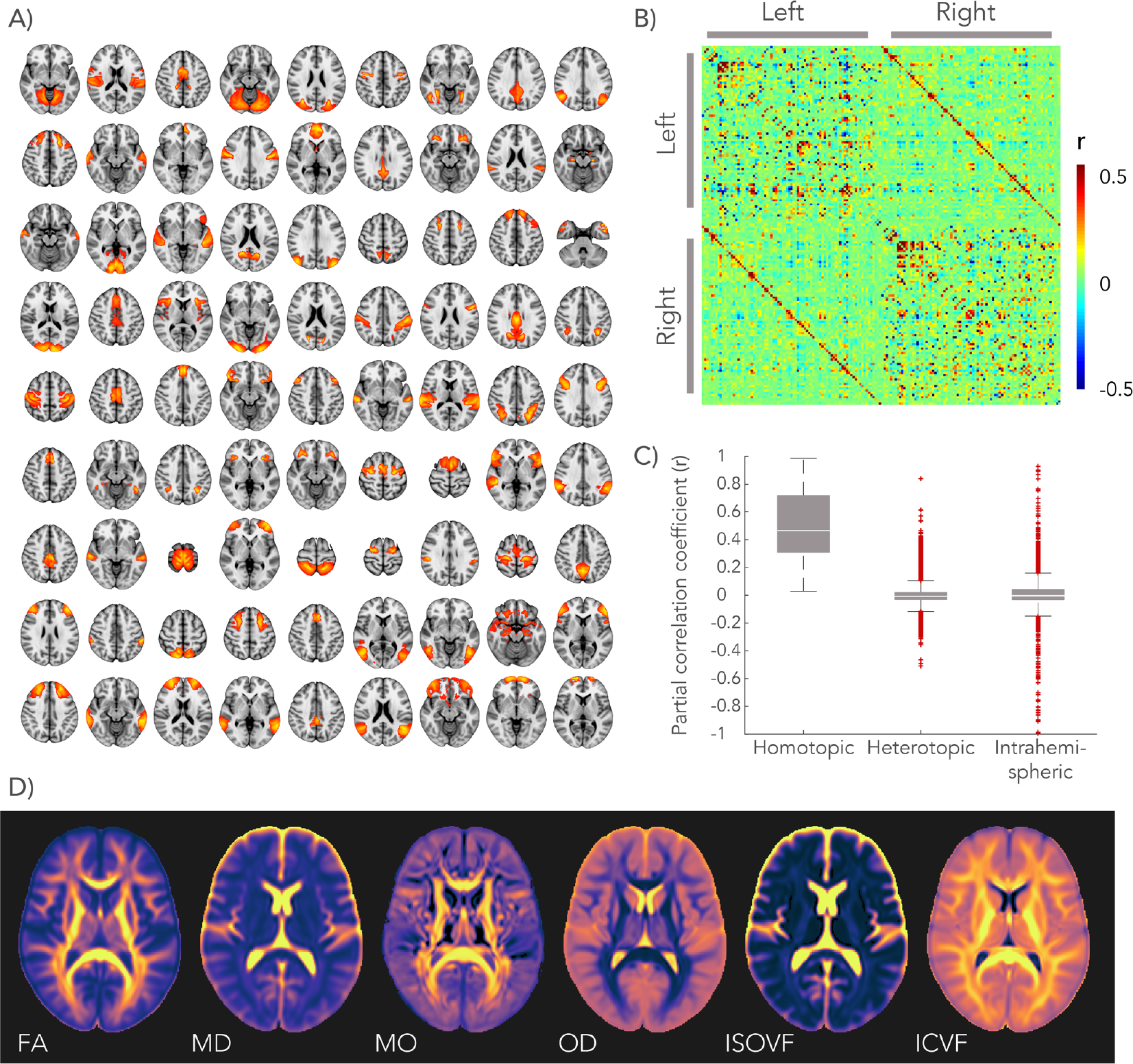
Functional parcellation of the brain and dMRI derived microstructural maps. **A)** Functional parcels were identified by applying independent component analysis (ICA) to the resting-state fMRI data, splitting between the hemispheres, and isolating contiguous regions. Parcels representing 81 homotopic regions were paired between the two hemispheres by eye. **B)** Connectivity between homotopic pairs was estimated by partial correlation of the average time-series of each parcel as shown in the connectome (matrix entries are sorted first by hemisphere and then by parcel number). **C)** Strength (partial correlation) of different functional connections in the brain, sorted by type. The centre line depicts the median correlation coefficient for a specific type of connection; box limits, the 25^th^ and 75^th^ percentiles of the correlation coefficients; the whiskers extend to the most extreme data points excluding outliers (marked with a + symbol). **D)** Group-averaged microstructure maps derived from the dMRI data.

### White matter microstructural signature

For the white matter pathway connecting each pair of homotopic gray matter regions, a range of microstructural features was derived from the dMRI data. The diffusion tensor model describes the 3D water displacement profile at each voxel using an ellipsoid^21^. We extracted maps of fractional anisotropy (FA), mean diffusivity (MD) and anisotropy mode (MO)^22^ from this tensor fit. NODDI^16^ is a more biologically motivated model that aims to decompose the diffusion signal into an intra-cellular volume fraction (ICVF) and an isotropic volume fraction (ISOVF), the latter representing interstitial and cerebrospinal fluids. In addition, NODDI estimates an Orientation Dispersion (OD) index that quantifies the spread of fibres within the intra-cellular compartment. These dMRI-derived metrics represent an average across thousands of cellular components within each imaging voxel (2×2×2 mm^3^), producing whole-brain maps that provide mesoscopic information about how these features vary between different tracts and along a given pathway. Fig. 1.D depicts a brain map of each microstructural metric averaged across all subjects. The white matter pathway that connects a given homotopic region pair was identified using probabilistic tractography^23^ performed on the dMRI data between the regions.

### Predicting functional connectivity with microstructure

We performed a multiple regression analysis to test whether dMRI microstructural features could predict cross-subject patterns of functional connectivity in the main cohort of 7,481 subjects. For a given homotopic pair of regions, the functional connectivity for all subjects was represented as a vector (N_subjects_ × 1). To model the microstructural signature, a corresponding matrix was constructed for the white matter pathway connecting the homotopic pair. Rather than averaging the microstructural profile along the tract, we created complete microstructural signatures that account for the spatial variability along the tract. These are matrices where each row contains, from a given subject, one or more microstructural parameters, each estimated in all voxels lying along the tract centre. These microstructure matrices are large (N_subjects_ × N_voxels_), resulting in too few degrees of freedom to robustly perform a direct regression (Nvoxels = 5750 (SD 4000)). Thus, a principal component analysis (PCA) was performed on each microstructural matrix, from which the top 30 principal components (see Supplementary Fig. 1) were extracted to serve as a set of regressors (N_subjects_ × 30) (see Fig. 2 for an overview). Seven different linear models were created for each homotopic pair: one for each of the dMRI-derived metrics (FA, MD, MO, OD, ISOVF, ICVF) and a multimodal approach combining all these microstructural metrics in a single matrix. For the multi-modal matrix, the microstructural matrix for each metric was first normalized by its first singular value, and these normalized matrices were concatenated to form a single multimodal matrix that was fed into the PCA.

**Figure 2.**
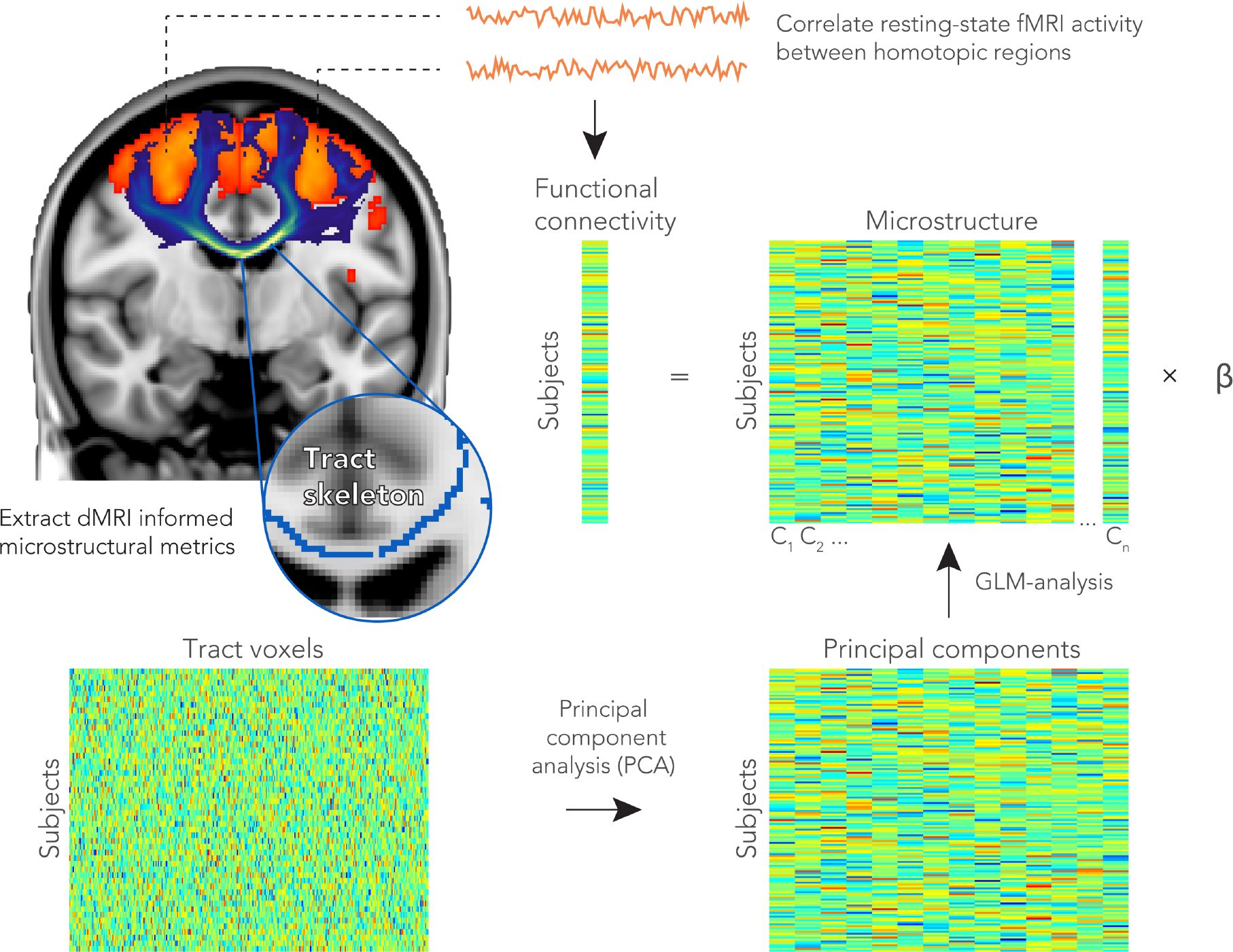
Prediction of functional homotopic connectivity from white matter microstructure. Between a pair of functionally defined homotopic areas (shown in orange in the brain), probabilistic tractography was performed to delineate the neuronal tract (shown in blue). The white matter skeleton voxels within a tract were stored as rows in a matrix for each subject. To extract the highest cross-subject variance among the TBSS voxels for a given microstructure metric, we performed a dimensionality reduction on this matrix using a principal components analysis. The top principal components (n = 30) were fed into a linear regression model as explanatory variables for the functional connectivity between a homotopic pair.

We assessed the statistical significance of the 30 regressors’ beta values in each model with permutation testing. Permutation testing was performed independently across the homotopic pairs and models (see Methods section). The significance (p<0.05, corrected for multiple comparisons) is indicated per microstructural metric in Fig. 3. For all microstructure metrics, the overall regression model was able to predict a statistically significant amount of cross-subject variance in most (68-86%, depending on choice of metrics) of the tested homotopic pairs, indicating a relationship between microstructure and functional connectivity. We can further consider individual regressors (i.e., specific principal components). The statistically significant beta values generally correspond to the top principal components (left-most columns in Fig. 3). This indicates that the highest cross-subject modes of microstructural variation also explain the most cross-subject variation in functional connectivity. As the regressors reflect the primary modes of variation in the dMRI data but are used to model the fMRI data, this property is not trivially guaranteed. For some regions, no significant associations were found between homotopic functional connectivity and a given microstructure metric. The multi-modal microstructure model combining across specific metrics resulted in the largest number of significant regressors, as well as providing an overall significant prediction of functional connectivity for the largest number of regions – 70, representing 86% of the total brain areas considered.

**Figure 3.**
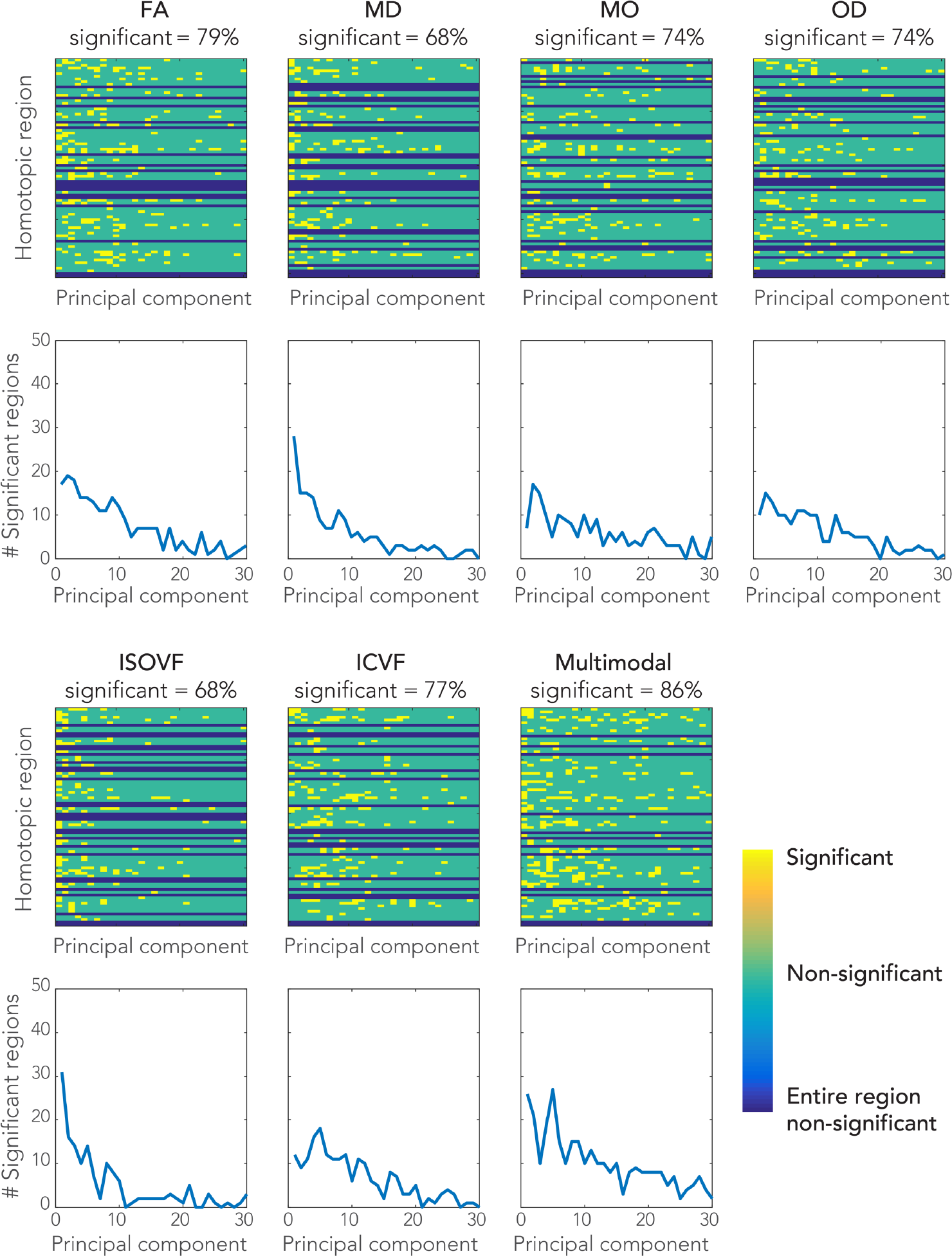
Significant associations between functional connectivity and microstructure of the connecting white matter tract. Each row in the matrices represents a homotopic region pair with each entry a regressor (on the microstructural principal components) of the linear model. Significance of the regressors is color-coded. The graphs depict the number of significant regressors found for each of the principal components. The percentage of homotopic region pairs demonstrating at least one significant regressor is given in the label of each matrix.

We additionally evaluated the regression models in terms of the total variance explained (TVE) in functional connectivity by the microstructural metrics. Substantial variation in TVE was found across the different brain regions investigated. For example, in the multimodal regression across all significant model fits, the minimum TVE was 0.73% (middle temporal gyrus) and the maximum TVE was 10.8% (posterior cingulate cortex). By mapping the TVE scores back to the 81 homotopic regions, we can visualize the spatial pattern of brain areas whose degree of functional connectivity was explained by the underlying microstructure (Fig. 5A). We also computed Z-scores to summarize the overall model fits. The multi-modal microstructure regression model yielded on average a higher score than the regressions with any single microstructural metric (Z = 11.5 Fig. 4), suggesting that the different microstructural metrics explain different variance in functional connectivity. The model incorporating FA shows the highest average Z-score of all individual metrics (Z = 9.6 ± 4.7), although the different metrics are overall fairly similar: ICVF (Z = 8.9), OD (Z = 8.8), MO (Z = 8.7), MD (Z = 8.3), ISOVF (Z = 8.1). A list of all brain areas investigated with their corresponding TVE values for the multimodal microstructure model is given in Supplementary Table 1.

**Figure 4.**
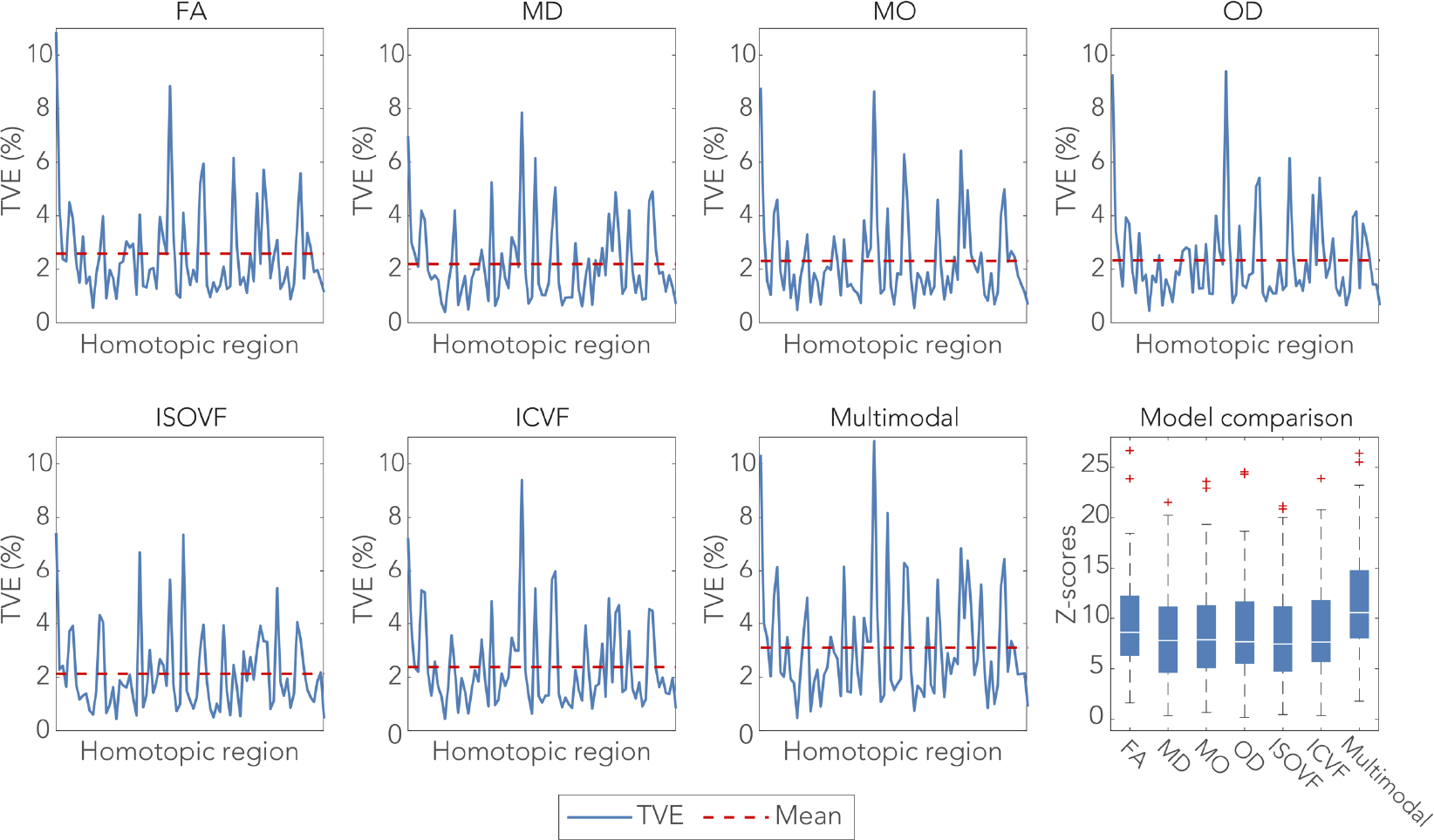
Total variance explained (TVE) in the functional connectivity of each homotopic region pair by the microstructural metrics derived from the connecting white matter tract. The box-and-whisker plots on the bottom right depicts the model performance of each metric in terms of an F-to-Z transformed score. The centre line depicts the median Z-scores across the homotopic region pairs; box limits, the 25^th^ and 75^th^ percentiles of the Z-scores; the whiskers extend to the most extreme data points excluding outliers (marked with a + symbol).

Tensor-based features (FA and MD in particular) have been shown to provide sensitive indicators of changes to tissue microstructure in a broad range of contexts. However, these measures can be influenced by multiple aspects of tissue microstructure^13^, making interpretation difficult. We tested whether functional connectivity relates to a microstructure feature with greater biological specificity. Here, we build on our previous work demonstrating this - quantitative agreement between fibre orientation dispersion estimates derived from dMRI data and dispersion estimated from structure tensor filtering of myelin stains in the same post-mortem human brain tissue^24^. The callosal fibre dispersion profile correlated well between the ex-vivo imaging data and the in-vivo dMRI NODDI analyses presented above, with both methods indicating high dispersion on the midline and lower dispersion in the lateral aspects of the callosum. Furthermore, fibre dispersion at the midline of the corpus callosum was able to explain significant variance in interhemispheric functional connectivity (Supplementary Fig. 2). While the explained variance was less than with the spatially-extended microstructure models presented above, the validation against histology demonstrates biological specificity of this particular association.

### Model validation

We further tested the validity of the above models by applying them to the replication cohort of 3,873 subjects. Because the models are applied directly and not retrained to fit to the new subjects, this constitutes a direct prediction of functional connectivity from dMRI data in unseen subjects. Each replication subject’s data was projected onto the 30 top principal components and then multiplied by the regression coefficients estimated from the original cohort to predict that subject’s functional connectivity. As shown in Fig. 5, the TVE was quantitatively very similar from region to region in the previously unseen subjects as in the main cohort upon which the model was based.

**Figure 5.**
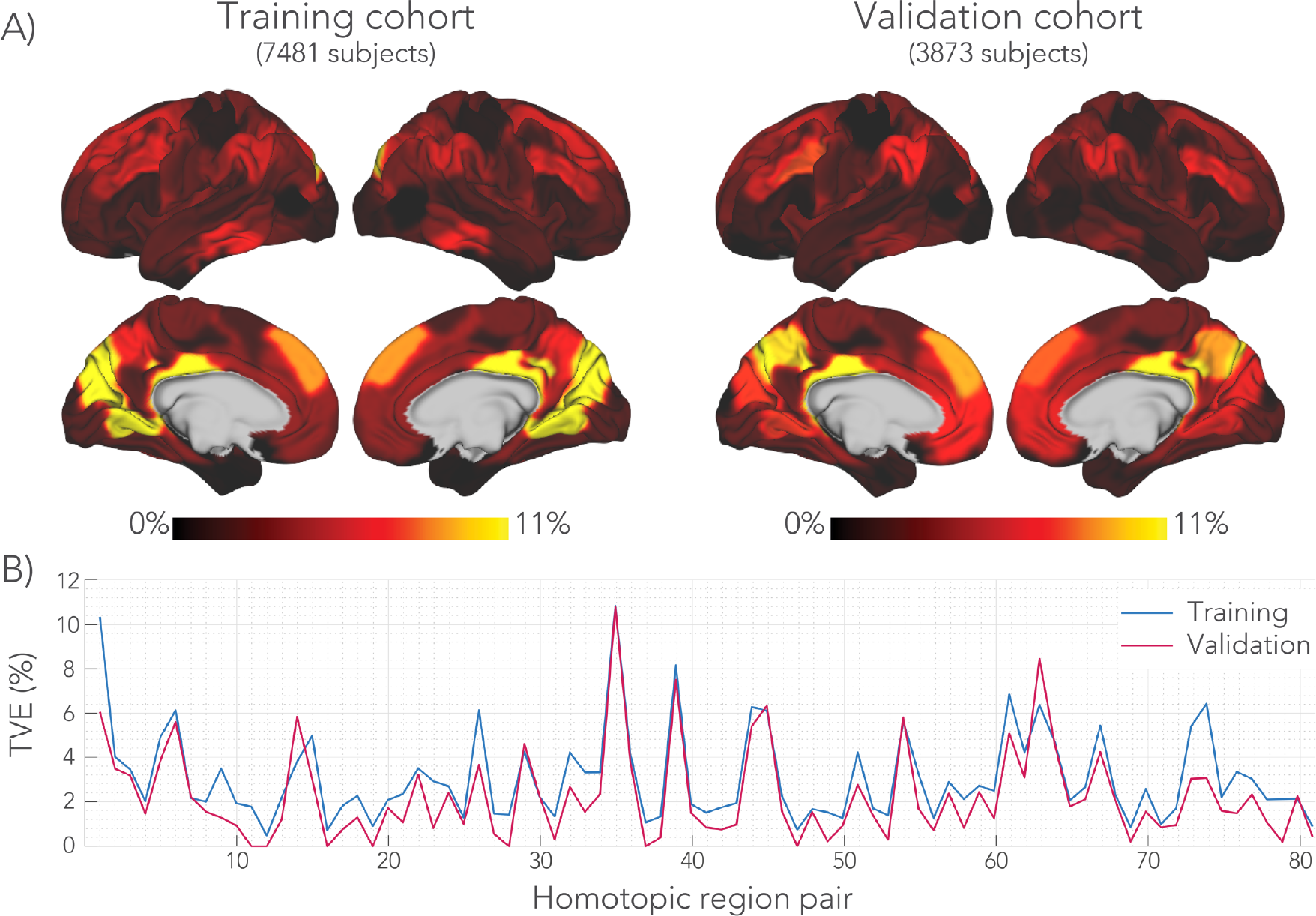
Total variance explained (TVE) by the multimodal regression model. **A)** TVE values mapped onto the brain surface for each of the homotopic regions. The maps were smoothed with 2 mm Gaussian kernel to aid visualization. A similar pattern across the brain was found for the regression models incorporating the individual microstructural metrics. **B)** Graph depicting the TVE values for each homotopic region. The model was trained on the main cohort of 7481 subjects. By applying the regression models without further fitting, we could predict functional connectivity in the additional validation cohort of 3873 unseen subjects. The homotopic region numbers correspond to the brain areas listed in Supplementary Table 1.

Several medial regions have a particularly high TVE score, with in particular the posterior cingulate cortex and the intra-calcarine cortex having TVE over 10%. Regions in the temporal lobe, ventral parts of frontal lobe and lateral aspect of the occipital lobe demonstrate the lowest TVE scores.

### Negative control analysis

Although the above analyses suggest a general microstructure-function relationship, it is not clear whether these associations are specific to the pathway connecting a given pair of regions, or whether functional connectivity reflects global variance in the microstructural metrics across subjects. A new series of regression analyses were performed similar to those depicted in Fig. 2, but instead of taking microstructure from the specific callosal pathway connecting a homotopic pair, the microstructure was derived from a different “wrong” tract (Fig. 6.A). From the 81 callosal sub-regions defined above, we selected a subset of 30 distinct tracts that shared minimal spatial overlap (Supplementary Fig. 3) for use as control (“wrong”) tracts. We then assessed whether any of the control tract regressions had similar performance to the correct tract (Fig. 6.C). For 64% of the homotopic areas, the highest Z-score was obtained when the model was performed with the anatomically correct tract; overall, for 81% of brain areas the correct tract ranked among the top three models (Fig. 6.D).

**Figure 6.**
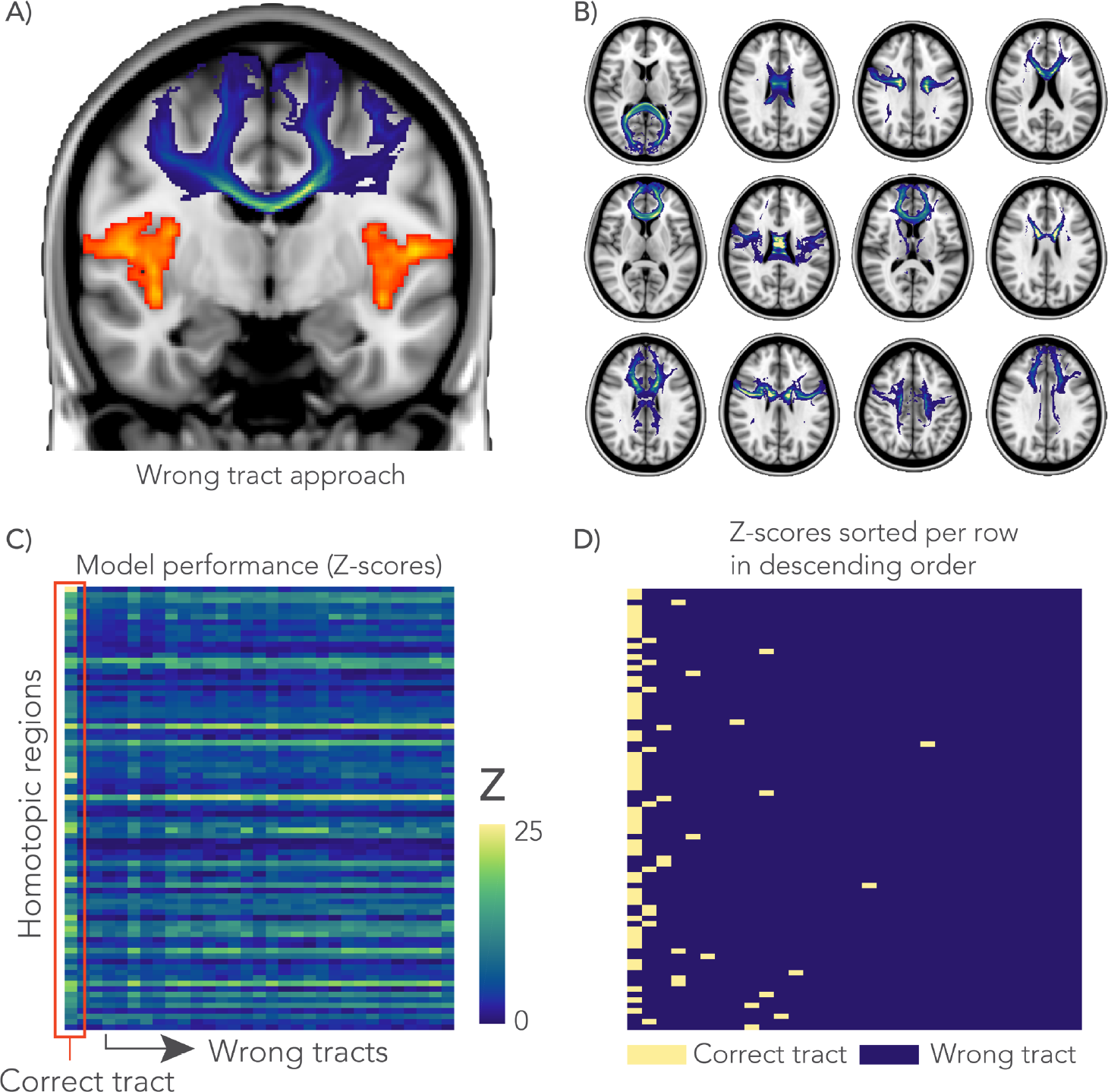
Negative control analysis. **A)** In the wrong tract approach, the GLM-analysis was performed with a tract (shown in blue) that does not directly connect between a homotopic pair of interest (shown in orange). **B)** A total of 30 distinct tracts (12 shown here) were chosen based on minimal spatial overlap between them. **C)** For each GLM, an F-statistic (across the whole model, with the degrees of freedom for model and error, 30 and 7450, respectively) was calculated and transformed to a Z-score to compare between the correct tract and 30 wrong tracts. All GLMs in this analysis were derived from the multimodal microstructural information. **D)** Rows from the matrix in C were sorted in descending order of Z-score and labelled according to whether they represent the correct pathway or a different tract. The highest Z-scores (left-most column) correspond to the anatomically correct tract in 64% of cases, and overall the correct tract was in the top three models in 81% of cases.

### Genome-wide associations

We studied the influence of genetics on the microstructure-function relationships reported in this work via a genome-wide association study (GWAS). All subjects in this analysis were selected based on recent British ancestry and availability of genotype data that passed the quality control procedures of UK Biobank^17^.

For the GWAS, we considered the model fits of the multimodal microstructure model that predicts functional connectivity between a pair of homotopic areas as the phenotype (see Methods section). The model fits were derived for all 81 homotopic brain region pairs (see Supplementary Fig. 4) and fed into the GWAS. The model fit of each homotopic region pair was evaluated against a total 11,734,353 single-nucleotide polymorphisms (SNPs). Figure 7 depicts the association across SNPs for the homotopic pair with the largest TVE in the multi-modal microstructure model (i.e., the posterior cingulate cortex). A group of SNPs in chromosome 14 demonstrated a strong association with the microstructure-function phenotype. These SNPs were co-located with the *DAAM1* gene (Dishevelled Associated Activator of Morphogenesis 1), and some were also within *DAAM1*’s promoter region (regulating expression of the gene)^25^. Expression of the *JKAMP* gene (Jun N-Terminal Kinase 1-Associated Membrane Protein) was also regulated by these SNPs, as demonstrated by 3D chromatin interaction data^26^, included in the Virtual 4C online resource^27^. *DAAM1* plays an important role in the Wnt signalling pathway inside the cell, indirectly regulating cell polarity and movement during development. In the central nervous system, this gene has been shown to facilitate the guidance of commissural axons at embryonic stage in mice and drosophila^28,29^. Furthermore, the GWAS revealed many SNPs within the *LPAR1* gene (Lysophosphatidic Acid Receptor 1) located in chromosome 9. *LPAR1* is one of the six receptors involved in the lysophosphatidic acid signaling pathway in the cell^30^. SNPs co-located with both *DAAM1* and *LPAR1* were found for the microstructure-function association of multiple brain areas (Fig. 7). Detailed Manhattan plots at the location of *LPAR1* and *DAAM1* are given in Supplementary Figures 5 and 6, respectively. A collection of SNPs was found for other regions of the brain, some of which relate to neural organization, metabolism or signalling (Table 1). Manhattan plots depicting the GWAS for the microstructure-function model fits of each homotopic region pair in the discovery cohort can be found in Supplementary Figure 8.

**Figure 7.**
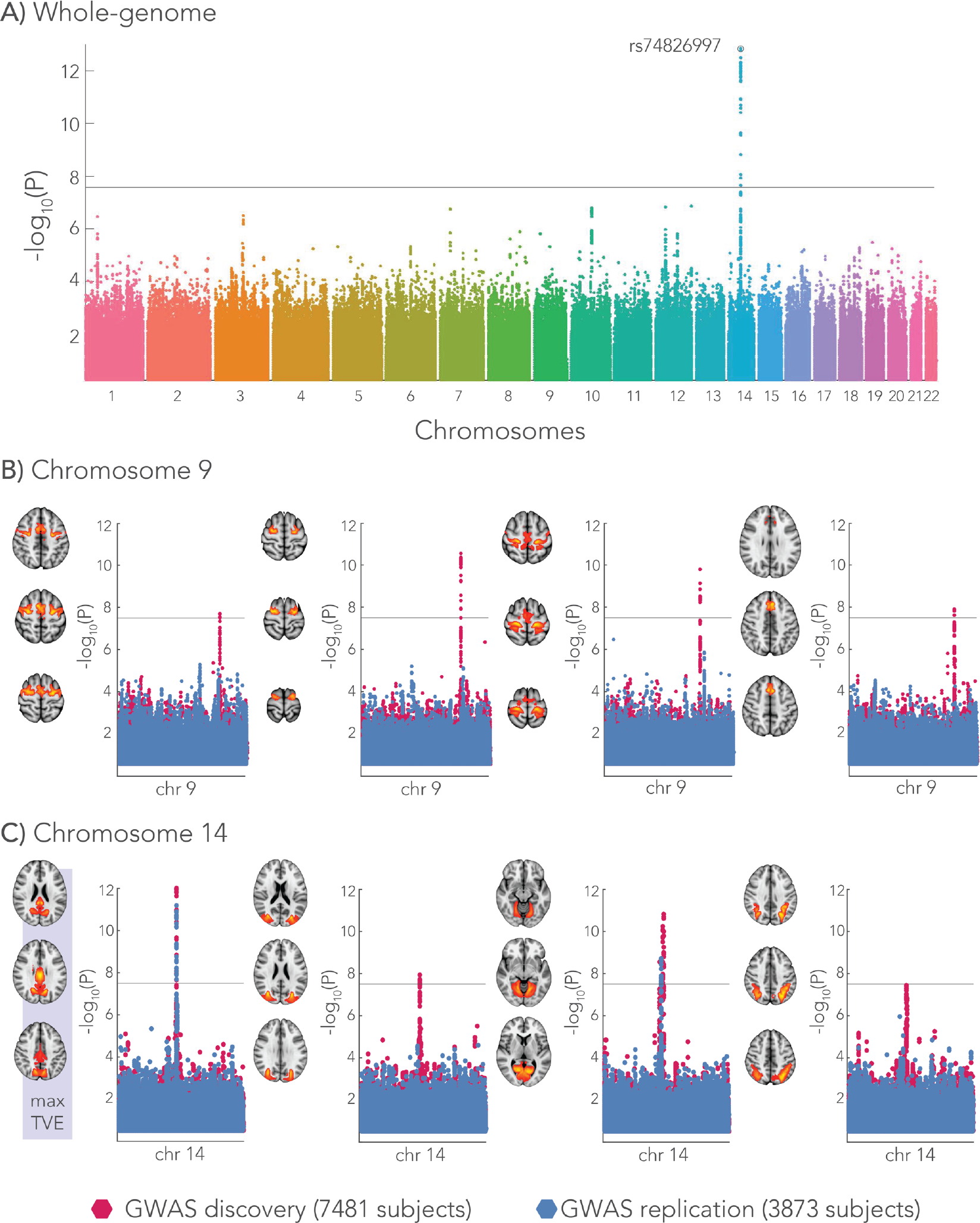
Genome-wide associations with the microstructure-function phenotype (i.e. the pattern of functional connectivity that can be predicted from white matter microstructure). The Manhattan plot depicts the associations with each SNP across all chromosomes expressed as the −log10 p-value. **A)** The genome-wide Manhattan plot is given for the homotopic brain area showing the highest total-variance-explained by the microstructure – function model and yields the strongest association with a SNP (rs74826997) in chromosome 14. Furthermore, single chromosome Manhattan plots are shown for a range of brain areas. The microstructure – function model fits of these areas were repeatedly associated with SNPs in either chromosome 9 **(B)** or 14 **(C)** that co-located with the genes *LPAR1* and *DAAM1*, respectively. The discovery GWAS was carried out with 7481 subjects revealing the group of SNPs (red dots in single chromosome Manhattan plots). In an additional cohort of 3873 subjects, we aimed to replicate these hits (blue dots in single chromosome Manhattan plots). The spatial maps of these brain areas are given for each of Manhattan plot. The brain area (posterior cingulate cortex) highlighted with *max TVE* corresponds to the genome-wide Manhattan plot at the top of the figure. A significance threshold is given for a −log10(p-value) equal to 7.5 corresponding to a p-value of ~3 × 10^−8^.

The GWAS was repeated for subjects in the replication cohort. Rather than using the model prediction approach described above, the multi-modal microstructure models were first re-trained to obtain a prediction of functional connectivity for these subjects. This approach was motivated by the lower TVE for the model predictions compared to the original fits (Fig. 5B), suggesting that in this smaller replication cohort, biases in the accuracy of the microstructure-function model could reduce sensitivity. Additionally, this approach makes the genetic replication analysis more fully independent of the discovery dataset. Replication GWAS was performed on microstructure-function phenotypes from the homotopic regions showing an association in chromosome 9 and 14 in the original subjects. SNPs associated with two out of three brain areas in chromosome 14 replicated. The SNPs in chromosome 9 within the *LPAR1* gene were not found in the replication GWAS (Fig. 7).

Thus far, the reported SNPs were found using microstructure-function model fits as the target phenotype. However, this result could simply reflect that both functional connectivity and microstructure correlate to these SNPs. To test for specificity, two additional GWAS were run using the functional connectivity and multimodal microstructure (1^st^ principal component) as the target phenotypes. Results are depicted for the homotopic region pair with the largest TVE for the model fit, given in Supplementary Figure 7. No SNPs co-located with either *DAAM1* or *LPAR1* were found in these GWAS for any homotopic regions. This suggests that the relationship to *DAAM1* and *LPAR1* is more specific to the specific component of functional connectivity that can be predicted by white matter microstructure. The GWAS associating with microstructure yielded SNPs within the *VCAN* gene, which were previously found to associate with ICVF throughout white matter in the brain^18^.

**Table 1.** Genome-wide associations with the microstructure-function phenotype (i.e. the pattern of functional connectivity that can be predicted from white matter microstructure). Listed are rsids of the SNPs showing the most significant association. Some SNPs were associated with the microstructure-function model fits of multiple homotopic region pairs (highlighted in gray). The nearest gene of each SNP is reported with its possible function in the human central nervous system. Furthermore, the base-pair position, the SNP alleles, minor allele frequency (maf) and the p-value of the discovery GWAS are given.

| chr | rsid | number of areas -<br>location in brain | nearest<br>gene | function in central nervous<br>system | position | ref<br>allele | minor<br>allele | maf | p-value |
| --- | --- | --- | --- | --- | --- | --- | --- | --- | --- |
| 1 | rs201286854 | 1 – Sensory-Motor | MACF1 | Links microtubuli with actin in<br>cytoskeleton of the cell | 39938494 | C | A | 0.011 | 2.23E-08 |
| 2 | rs7582436 | 1 – Medial occipital | ATL2 | Unknown | 38532584 | A | G | 0.371 | 2.88E-09 |
| 6 | rs12200595 | 1 – Occipital | NHSL1 | Associated with Nance-Horan<br>syndrome | 138862574 | G | T | 0.277 | 9.12E-09 |
| 7 | rs7788173 | 1 – Frontal pole | SEM1 | Unknown | 96219460 | A | G | 0.689 | 3.09E-09 |
| 7 | rs143137457 | 1 – Superior<br>temporal | CUX1 | Regulates dendritic branching,<br>spine morphology and synapses | 101425857 | G | A | 0.012 | 1.07E-08 |
| 9 | rs4556147 | 4 – Prefrontal | LPAR1 | Lysophosphatidic Acid Signaling<br>in central nervous system | 113651161 | A | T | 0.221 | 2.82E-11 |
| 14 | rs74826997 | 3 – Medial occipital | DAAM1 | Wnt signaling pathway, axonal<br>growth and guidance | 59628609 | T | C | 0.125 | 1.41E-13 |
| 15 | rs4924345 | 1 – Lateral prefrontal | C15orf54 | Associated with spinal cord<br>tumor | 39639898 | A | C | 0.081 | 8.32E-09 |
| 19 | rs117042631 | 1 – Middle frontal | PODNL1 | Unknown | 14053839 | G | A | 0.033 | 1.10E-08 |
| 20 | rs4911180 | 1 – Frontal orbital | UQCC1 | Unknown | 33972948 | G | A | 0.626 | 1.82E-08 |

## Discussion

Although basic principles relating axonal properties to neural signalling are well established, the degree to which functional connectivity is mediated by microstructural organization at the level of macroscopic tracts is largely unknown. Several studies have related the “strength” and topology of structural connections to functional activity based on fMRI and dMRI^10,31^, but these studies are uninformative about microstructure. Here we focused on commissural fibres through the corpus callosum, a set of connections which can be readily measured using MRI both structurally and functionally. Our results are consistent with previous work^19,20^ in that connections between homotopic areas in both hemispheres were functionally the strongest connections in the brain. Evidence that functional interhemispheric connectivity is indeed primarily facilitated by axons running through the corpus callosum comes from callosotomy studies (i.e., surgical sectioning of the corpus callosum) in both animals^32^ and humans^33^.

We have demonstrated that white matter microstructure is strongly associated with functional connectivity. Replication in nearly 4000 subjects demonstrates that the regression models fit in the main cohort have predictive power in unseen subjects, including good reproducibility in terms of the total amount of variance of functional connectivity explained in a given region. In both the main – and validation cohort, functional connectivity variance was best explained in brain region close to the medial aspect of the brain, for example the intra-calcarine – and posterior cingulate cortex (see Fig. 5). It should be acknowledged that some of these regions emerged as single contiguous node after spatial ICA as opposed to more distal homotopic pairs separated by other brain structures (see Fig. 1). However, it is unclear why such functional principle would be better predicted by a completely independent measure of white matter microstructure estimated with diffusion MRI. The hypothesis that the explained variance was higher if there is little white matter separating a homotopic pair, was not supported by a simple correlation of the number of white matter voxels in each model with the TVE across homotopic region pairs (r=−0.17, p-value=0.13). Moreover, the SNPs found reported by the GWAS do not necessarily correlate solely with microstructure-function model fits of homotopic pairs close to the midline of the brain, suggesting that these fits are biologically informative.

For the majority of homotopic area-pairs connected via the corpus callosum, the strongest model prediction was derived from microstructure in the anatomically-correct pathway, compared to microstructure obtained from any of a large number of other callosal pathways. This negative control analysis is informative because it establishes that microstructure-function relationships have a high degree of regional specificity and do not simply reflect global (brain-wide) inter-individual differences in microstructure and associated function. A similar result related FA in the cingulum to resting-state functional connectivity between posterior cingulate and medial-frontal cortices^14^. Interestingly, for a minority of the brain areas investigated, functional connectivity was better explained by microstructure from another white matter tract. This is primarily true for frontotemporal regions, where on average lower functional homotopic connectivity was found, in agreement with previous literature^34^. These regions may not receive much callosal input and are primarily connected to intra-hemispheric brain areas via associations fibres^35^. Also, temporo-polar regions connect more likely via the anterior commissure, a tract that was not included in our analyses.

Imaging microstructure with dMRI is a rapidly evolving field, including many models that were only recently developed. Though the microstructure metrics are sensitive to tissue architecture at microscopic level, evaluation against reference measures such as histology is essential. As such, we demonstrated good correspondence between orientation dispersion (OD) profiles derived from the corpus callosum in ex-vivo dMRI and myelin staining^24^, providing confidence in the biological meaning of this specific measure. In agreement with histology^36,37^, the dMRI data used in our study indicates that fibres are more dispersed at the centre of the corpus callosum as compared to its lateral aspects. Here, we demonstrate that this validated measure of fibre dispersion also relates to interhemispheric connectivity of some homotopic areas (Supplementary Fig. 2), albeit with less explanatory power than our regression models that incorporate the full spatial richness of microstructure metrics across the white matter tract. Furthermore, the biological interpretation of tensor-derived measures is less clear and would require further investigation.

Richness of the data in the UK Biobank project allowed us to associate genetic variants with the imaging derived phenotypes in this study. While investigations of genetic influences on brain structure and function were previously mostly limited to specific candidate genes (for example, the involvement of the ApoE gene in Alzheimer’s disease^38^), GWA studies evaluate all common genetic variants against a certain phenotype ^39^. We conducted a GWAS associating SNPs with the fraction of functional connectivity that was predicted by microstructure, for each homotopic region pair. In chromosomes 9 and 14, a group of SNPs was found showing a strong association with the cross-subject pattern of functional connectivity predicted by microstructure for multiple brain areas (Fig. 7). As these SNPs were not found in a GWAS relating solely functional connectivity or microstructure, these SNPs appear to be unique to the microstructure – function relationship (see Supplementary Fig. 7). However, previous work relating cortical thickness to genetic variants also reported SNPs co-located with *DAAM1* in the cuneus area^18^ (http://big.stats.ox.ac.uk). For the replication cohort, the SNPs in chromosome 14 – co-located with genes *DAAM1* and *JKAMP* – were only replicated for two of the three brain areas showing hits in the discovery GWAS. We could not reproduce the SNPs in chromosome 9 – within the *LPAR1* gene – for the brain areas previously showing these associations. If these relationships do exist, it may simply replication may have failed. First, the number of subjects in the replication cohort is almost half of the number in the discovery cohort and thus suffers from lower statistical power. Indeed, the SNPs in chromosome 9 express p-values that are just above the significance threshold in the discovery GWAS (Fig. 7). Second, the microstructure - function model fits that were fed into the GWAS demonstrate varying performance in terms of the TVE between the main and replication cohort. Through refitting the models aim to predict functional connectivity in a similar manner based upon white matter microstructure (Fig. 5), they may explain a different component of variance regardless of the sample size.

The identified SNPs in chromosomes 9 and 14 have previously been shown to be important for brain development. The *DAAM1* gene is expressed in many tissue of the human body and plays an important role in the Wnt signalling pathway^40^. In neuronal tissue, *DAAM1* is primarily found in the shaft of neuronal dendrites^41^ and in the developing brain it aids axonal guidance in targeting distal brain regions^42^. Knock-out studies in mice and drosophila have shown deficits in the central nervous system when *DAAM1* is not expressed^28^. In particular, the formation of commissural fibres at an embryonic stage was disturbed^29^. 3D chromatin data revealed that the SNPs in chromosome 14 also regulate expression of the *JKAMP* gene^26^. While diseases associated with *JKAMP* include medulloblastomas^43^, its exact mechanism in brain development is not well described in literature. For chromosome 9, several SNPs were located in the *LPAR1* gene, a receptor involved in the lysophosphatidic acid signalling pathway. These receptors are found on the membranes of most cell types in the central nervous system and have been linked to some neural processes including but not limited to neurogenesis, myelination, microglial activation, and astrocytes responses^30,44^.

The degree to which functional connectivity between brain regions is mediated by microscopic properties (microstructure) of the white matter pathways is a fundamental question in neuroscience. We demonstrated that a large fraction of variation in inter-hemispheric functional connectivity can be predicted from white matter tract microstructure connecting two homotopic regions. Our results suggest that microstructure-function relationships are general (across many brain regions), specific (based on analysis of control tracts) and reproducible (in a replication cohort). Furthermore, the microstructure-function association was underpinned by genetic variants and in particular with SNPs co-located with the genes *DAAM1* and *LPAR1*, identified in multiple brain regions. To conclude, genetically-determined properties of white matter microstructure sculpt brain activity. Attribution of these relationships to specific biological sources - and ideally causality - cannot be achieved with this kind of observational study but would likely require interventional studies in animals.

## Materials & Methods

### Data acquisition and pre-processing

We used resting-state functional MRI and diffusion MRI data provided by the UK Biobank project. An extensive overview of the data acquisition protocols and image processing carried out on behalf of UK Biobank can be found elsewhere^7,45^. Description of post-processing pipelines and acquisition protocols of MRI data in UK Biobank are available at http://biobank.ctsu.ox.ac.uk/crystal/docs/brain_mri.pdf. Unless stated otherwise, processing of the MR images was performed using FSL^46^. All imaging data was acquired on a 3T Siemens Skyra MRI scanner (software platform VD13) using a 32-channel receive head coil.

Resting-state fMRI data with 2.4 mm isotropic resolution and whole-brain coverage (field of view, 88×88×64 matrix) was acquired in a six-minute session (multiband acceleration 8, TR=0.735 ms, 490 time-points). The functional data was motion corrected ^47^ and FIX-cleaned^48^ to remove physiological noise and image artefacts, before transforming the data to a 2 mm MNI-template.

Diffusion MRI data were acquired at 2 mm isotropic resolution achieving whole brain coverage (field of view, 104×104×72 matrix) with two b-values (b=1000, 2000s/mm^2^), with 100 unique gradient directions over the two shells (50 directions/shell). with a voxel size of 2 mm isotropic. The total acquisition time was 6.5 minutes (multi-band acceleration 3, TE/TR was 92/3600 ms). After eddy current correction of all images^49^, tensor metrics (FA, MD, MO) were calculated from the lower shell (b=1000s/mm^2^) using DTIFIT. Both shells were used to estimate the NODDI model metrics (ICVF, ISOVF, OD) using the AMICO toolbox^50^. All microstructural metrics were projected onto a white matter skeleton using TBSS (Tract-Based Spatial Statistics^51^) to minimize misalignment of tracts due to inter-subject morphology and registration errors.

### fMRI processing

The resting-state fMRI data were fed into an Independent Component Analysis (ICA) using the MELODIC tool^52^ to identify resting-state networks present on average in the whole population. First, data was reduced to 100 dimensions using PCA and then fed into spatial ICA, from which 55 components corresponded to functional regions, and the other 45 judged to reflect physiological noise or image artifacts (“noise”). A functional component was split if it consisted of non-contiguous brain regions, yielding 81 bilateral (homotopic) regions that were further split between the hemispheres to estimate interhemispheric connectivity (see Supplementary Table 1). Average time-series were generated for all ICA components (i.e., homotopic areas and noise components) by a spatial regression of the subject’s voxelwise resting-state fMRI time-series with the ICA spatial maps. Further analyses were performed using the FSLNets toolbox (https://fsl.fmrib.ox.ac.uk/fsl/fslwiki/FSLNets). The average time-series within a homotopic area were demeaned and “cleaned” by regressing out the time-series from the 45 noise time courses. Functional connectivity was estimated between all pairs of components (2×81) by means of partial correlation of the cleaned time-series using Ridge regression with a regularization factor ρ=1. Partial correlation aims to measure direct connectivity between two areas by first regressing out all other regions’ time-series before calculating the correlation (i.e., established through inversion of the covariance matrix).

### dMRI tractography

White matter tracts between functional regions were delineated using tractography. Up to three fibre orientations were fitted at each dMRI voxel in a Bayesian approach using bedpostX^53^ modified for multi-shell data^54^. Probabilistic tractography was then performed with the probtrackx2 algorithm^23^ by generating streamlines from a seed region (5000/voxel) in one hemisphere and only saving streamlines that passed through the corpus callosum and terminated in the same region in the contralateral hemisphere. This process was repeated by switching the seed and the target area between the hemispheres. The overlap of the identified tracts in this two-step approach were used for further analysis. The tracts were generated for all 81 homotopic pairs (each representing either the seed or the target area) for 10 subjects drawn from the UK Biobank dataset. Tracts between a given homotopic pair were then averaged across these subjects and served as a tract for all subjects stored in 1 mm MNI-space, which was then further masked by the white matter skeleton voxels (derived from TBSS). Microstructural features derived from the tensor and NODDI fits were extracted from this final tract mask.

### Predicting functional connectivity from white matter microstructure

We used a multiple linear regression model to predict homotopic functional connectivity from a set of regressors describing the spatial pattern of microstructure along a white matter tract. The regression model was constructed for each pair of homotopic regions separately:

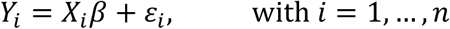

Here *Y_i_* (N_subjects_ × 1) is a vector that contains the functional connectivity values of all subjects derived from homotopic region *i* (over n = 81 regions). To build a model using *p* microstructural regressors, we need to estimate a set of regression coefficients *β* (*p* × 1) that describe the relative contribution from the microstructural metrics *X_i_* (N_subjects_ × *p*) along the white matter tract.

The regressors are derived in two stages. First, the microstructural metrics were extracted from the TBSS-voxels (white matter skeleton) corresponding to the tract of interest for every subject, yielding a matrix 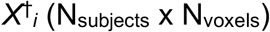. As the matrix 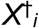 is very large, a direct regression with functional connectivity is ill conditioned. We therefore perform a dimensionality reduction on 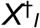 to derive a set of regressors reflecting the primary modes of variation of a given microstructural metric across space for the cohort of subjects. A singular value decomposition (SVD) was computed from matrix 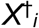, from which the top *p* components were retained, yielding matrix *X_i_* (N_subjects_ × *p*). In practice, *p* was set to 30 principal components, which approximately corresponded to a transition in the spectrum of singular values in terms of variance explained, above which variance explained roughly tracked noise singular vectors (Supplementary Fig. 1).

Matrices *X_i_* were constructed for each of the microstructure metrics separately, yielding six single-metric linear regression models per homotopic region. In addition, a multimodal regression model was created that combined across all microstructure metrics. For the multimodal regression, all raw microstructure matrices 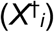 were normalized through division by their first singular value to ensure comparable range of values. The six normalized matrices were then concatenated and an SVD was performed on the concatenated matrix to reduce back to the top 30 components.

Finally, we defined a set of confound variables of no interest (*age, age*^2, *sex, age*sex, age*^2\**sex, resting-state fMRI head motion, and head size*) that could correlate with estimated microstructural measures (e.g. through artefacts such as partial volume) and thereby bias the estimated regressors. The confound variables were regressed out of the functional and microstructural data before fitting each regression model.

### Statistical analysis

Statistical significance of the regression models was assessed by means of permutation testing. A null distribution was constructed for each regressor by randomly permuting the functional connectivity values (the number of permutations was set to 100,000). A p-value (two-sided) was then determined in the non-permuted model from the null distribution. Because multiple models were evaluated, we corrected for the family wise error as in^55^. Here, we generated a maximum t-statistic distribution across all homotopic region pairs and regressors (i.e., the microstructural principal components) of the permuted t-statistics. From this maximum t-statistics null-distribution a corrected p-value was estimated for each of the non-permuted t-statistics. Furthermore, an F-statistic was computed to judge the overall performance of each regression model (degrees of freedom model and error, 30 and 7450, respectively). Finally, the effect size of the regression models was expressed in terms of total variance explained (TVE, similar to R-squared), describing the strength of the relationship between microstructure and functional connectivity.

### Negative control analysis

The statistical tests described above test whether there is a relationship between functional connectivity in a given brain region and the microstructure in the white matter pathway that connects them. However, this does not provide any insight into whether these relationships are specific: for example, microstructure and function could correlate at the whole-brain level. In this case, a regression model could indicate a statistically meaningful relationship even when using a white matter pathway that does not connect a given homotopic pair. Such a relationship could still be biologically meaningful, but the interpretation would change (e.g., individual brains could vary globally from hypo- to hyper-connected).

Accordingly, a negative control analysis was performed to evaluate the uniqueness of the microstructure-function relationships. From the 81 tracts in our study, a subset of 30 minimally overlapping tracts were selected as control (“wrong”) tracts. First, the Dice similarity index was computed among all tracts to quantify spatial overlap. Using k-means clustering (k=3 clusters), a cluster of tracts with the lowest average similarity indices was selected (Supplementary Fig 3).

The regression models were then re-evaluated for each homotopic area using the control tracts, rather than microstructure from the anatomically correct tract, i.e., the tract connecting the homotopic pair of interest. If, for a homotopic area, the anatomically correct tract was among the control tracts, an additional control tract was selected. To summarize, the regression models of the homotopic regions were performed once for the correct tract and 30 times with microstructure from the control tracts. Comparison between the correct and control tract analyses was conducted using the F-statistic.

### UK Biobank genetics data

The GWAS was performed using the BGENIE software^17^. Acquisition and processing steps of the genetics dataset for all subjects in the UK Biobank project can be found in^17^. For the discovery cohort, we began with the set of 12,623 brain imaged UK Biobank subjects for whom data were released in July 2017. As in (Elliott et al., 2017), to avoid confounding effects that may arise from population structure or environmental effects, we selected a subset of 8,522 unrelated subjects with recent British ancestry. Ancestry was determined using sample quality control information provided by UK Biobank^17^. We then filtered the genetic data to remove SNPs with minor allele frequency < 0.01% or a Hardy-Weinberg equilibrium p-value of less than 10^−7^, yielding a total of 11,734,353 SNPs distributed across the 22 autosomes. Not all of the UK Biobank subjects who underwent brain imaging have usable data with a given MRI modality. Of the 8,522 unrelated samples, we used a subset of 7,481 subjects which had usable dMRI and fMRI data according to previous quality control^45^. For the replication cohort, we examined a further set of 4,588 UK Biobank subjects for whom data were released by UK Biobank in January 2018. Application of the same inclusion criteria yielded 3,873 subjects.

### Ex-vivo MRI and histology data

MRI and microscopy data from three ex-vivo corpus callosum specimens were acquired and processed as described previously^24^. In brief, formalin fixed human brain tissue sections were scanned on a preclinical 9.4 T Varian MRI system. Diffusion MRI was performed with a spin-echo sequence with TE = 29 ms and TR = 2.4 s. Two shells were acquired (b = 2500 s/mm^2^ and b = 5000 s/mm^2^), each with 120 gradient directions and 0.4 mm isotropic resolution. Eight images with no diffusion weighting were acquired. A parametric model was fit to the dMRI signals from the b = 5000 s/mm^2^ dataset to obtain orientation dispersion (OD) estimates^56^.

Following MR scanning, the specimens were histologically sectioned and immunohistochemically stained for myelin (proteo-lipid-protein). The sections were digitized and we obtained fibre orientation estimates at each pixel using structure tensor analysis^57^. From a 2D local neighbourhood (0.4 × 0.4 mm) corresponding to the size of an MRI voxel, a fibre orientation distribution was computed from which orientation dispersion (OD) was derived. After registration of dMRI and microscopy data to the same image space^58^, dispersion estimates were compared against each other in the corpus callosum.

### Ethics and informed consent

All participants in the UK Biobank project signed an informed consent which is controlled by a dedicated Ethics and Guidance Council (http://www.ukbiobank.ac.uk/ethics). The Ethics and Governance Framework can be found at http://www.ukbiobank.ac.uk/wp-content/uploads/2011/05/EGF20082.pdf. IRB approval, also from the North West Multi-center Research Ethics Committee, was obtained for the Ethics and Governance Framework.

## Acknowledgements

All data in this study were obtained from the UK Biobank project (access number 8107). We are very grateful to all individuals who donated their time to participate in the UK Biobank study. K.L.M., M.K. and J.Mo. are supported by the Wellcome Trust (091509/Z/10/Z, 202788/Z/16/Z, 098369/Z/12/Z). The authors gratefully acknowledge funding from the Wellcome Trust UK Strategic Award [098369/Z/12/Z]. UK Biobank brain imaging and F.A.-A. are funded by the UK Medical Research Council and the Wellcome Trust. J.Ma. acknowledges funding for this work from the European Research Council (ERC; grant 617306) and the Leverhulme Trust. S.J. is supported by the UK Medical Research Council (MR/L009013/1). The Wellcome Centre for Integrative Neuroimaging is supported by core funding from the Wellcome Trust (203139/Z/16/Z). Finally, we would like to thank David G. Norris and Jose P. Marques for their valuable input on this work.

## Author contributions

J.Mo., S.M.S., S.J and K.L.M. designed the research. J.Mo. performed the research. K.L.M., F.A.-A., and S.M.S, developed acquisition and processing pipelines for the MRI data. L.T.E. and J.Ma. processed genetics data, provided tools for genome-wide associations analysis and gave feedback on genetics results. J.Mo., S.M.S., M.K., M.H., A.M.C.W., S.J and K.L.M. analysed the data provided useful interpretations in its outcomes. J.Mo. and K.L.M. wrote the manuscript, which was edited by all authors.

